# Pharmacokinetics of Sustained-release Buprenorphine and Extended-release Buprenorphine in Mice with Surgical Catheterization

**DOI:** 10.1101/2022.03.14.484329

**Authors:** Marissa Saenz, Elizabeth Bloom-Saldana, Tim Synold, Richard W Ermel, Patrick T Fueger, James B Finlay

## Abstract

*The Guide for the Care and Use of Laboratory Animals* strongly encourages the use of pharmaceutical grade chemicals and analgesics. The extra-label use of sustained-release buprenorphine (SRB) is commonly administered to rodents to mitigate moderate-to-severe pain. An FDA-indexed buprenorphine formulation, known as extended-release buprenorphine (XRB), has recently become available and is currently the only pharmaceutical grade slow-release buprenorphine approved for use in mice and rats. However, no studies have directly compared the pharmacokinetic (PK) parameters and therapeutic efficacy of SRB and XRB in surgically catheterized mice. Thus, we compared the plasma buprenorphine concentrations and PK parameters of SRB and XRB in mice after surgical catheterization. We hypothesized that mice treated with SRB or XRB would have circulating buprenorphine concentrations exceeding the therapeutic threshold for up to 72-hours post-operatively. Male and female C57Bl/6J mice were anesthetized, treated with either SRB (1 mg/kg, SC, once) or XRB (3.25 mg/kg, SC, once) and underwent surgical catheterization. At 6, 24, 48, and 72 h after SRB or XRB administration, arterial blood samples were collected. Post-operative weight loss was similar between groups with a decline of 11.7 ± 1.6 and 12.3 ± 0.7% in males and 7.6 ± 2.2 and 8.1 ± 1.1% (mean ± SEM) in females treated with SRB and XRB, respectively. Both SRB and XRB maintained circulating buprenorphine concentrations above the therapeutic level of 1.0 ng/mL for 72 h after administration. XRB buprenorphine concentrations were significantly greater (3-4-fold) than SRB concentrations at 6, 24, and 48 h, commensurate with the increase dose concentration of XRB to SRB. These results support the use of either SRB or XRB for the alleviation of postoperative pain in mice. The new availability of FDA-indexed XRB increases the options for safe and effective pharmaceutical grade analgesia in rodents.

## Introduction

Painful surgical procedures are often a major component of biomedical research and are commonly performed in laboratory animals.^1,2,11,16,23,32^ The AALAS position statement on *Alleviating Pain and Distress in Laboratory Animals* states that, “The avoidance and minimization of pain and distress in laboratory animals is an *ethical obligation* that preserves the welfare of animals used in research…”.^1^ Pain relieving measures often include adequate personnel training, environmental modifications, social housing, acclimatization to stressful procedures, and the use of anesthetic and analgesic drugs.^1,2,16,26^ Analgesic drugs are often one of the first tools used for the alleviation of painful surgical procedures because of their ability to decrease the anesthetic dose^28^ and prevent the development of central sensitization, also known as wind-up phenomenon.^11,23,32^ Furthermore, according to *The Guide for the Care and Use of Laboratory Animals*, “Successful surgical outcomes require appropriate attention to…[the] use of analgesics… during all phases of a protocol involving surgery and postoperative care”.^2^ Therefore, it is essential to alleviate undue pain and distress from otherwise painful surgical procedures using analgesics.

One of the most common classes of systemic analgesics used in laboratory rodents are opioids.^27^ Opioids have become the analgesic of choice in mice and rats because of their high therapeutic index^30^, minimal side-effects when used appropriately^22^, and effectiveness in alleviating moderate-to-severe pain.^4,9,11,17,19,32^ Buprenorphine is a partial μ-opioid receptor agonist and has been documented to be safe and effective in alleviating post-surgical pain in rodents.^4,9,11,17,19,25,27,30^ There are currently 3 formulations of buprenorphine that are commonly being used in laboratory rodents: buprenorphine HCl (BUP), sustained-release buprenorphine (SRB), and extended-release buprenorphine (XRB). The short duration of action of BUP presents a major limitation to its use^9^ and requires repeated handling and dosing every 3 to 12 hours.^12,18,21^ Alternatively, the slow-release buprenorphine formulations, SRB and XRB, have been reported to provide up to 72-hours of analgesia in mice after just one subcutaneous injection.^9,12,19,21,22,30^ Newly available XRB is FDA-indexed and the only current pharmaceutical grade slow-release buprenorphine for use in mice and rats.^19,30^ SRB is not considered pharmaceutical grade because SRB is a patented formulation produced in a commercial compounding pharmacy that can have variations in safety and efficacy based on the batches that are produced; however, many studies have independently evaluated SRB demonstrating its analgesic efficacy and safety in rodents.^4,11,12,17,21,22,35^ *The Guide* strongly encourages the use of pharmaceutical grade chemicals to ensure that unwanted side-effects are not introduced to a study^2^; thus, many institutions may consider switching from SRB to the newly available XRB. Yet, no studies have directly compared the pharmacokinetic (PK) parameters and therapeutic efficacy of SRB and XRB in mice with surgical catheterization.

Murine PK studies typically require terminal blood collections at various time points after drug administration to obtain adequate amounts of blood for analysis.^14^ However, blood sampling from different mice at various timepoints can introduce variability in inter-animal drug responses, as well as increase the number of animals required to complete a study.^14,29^ Ideally, permanent catheterization and repeated blood sampling in the same mouse would minimize interanimal variability and enhance the reproducibility of pre-clinical drug trial results.^6,7,29^ Furthermore, permanent carotid catheterization in mice has not resulted in changes in behavioral parameters, physiological parameters, and no detectable implication on animal welfare.^15^ Using carotid catheters to assess PK parameters and therapeutic drug efficacy is a refinement that decreases the number of animals used and decreases any variations in drug response due to animal-to-animal differences.^29^ Thus, the aim of this study was to compare the PK parameters and therapeutic efficacy of SRB and XRB in mice after permanent carotid artery and jugular vein catheterization. We hypothesized that mice treated with either SRB or XRB would have circulating buprenorphine concentrations exceeding the therapeutic threshold for up to 72-hours post-operatively.

## Materials and Methods

All animal procedures were performed in an AAALAC International-accredited facility and approved by the City of Hope Institutional Animal Care and Use Committee.

### Animals

Male (n=15) and female (n=12) 6-week-old C57BL/6J mice were purchased from Jackson Laboratory (Bar Harbor, ME) and allowed to grow and acclimate in the vivarium until catheterization surgery at 13-15 weeks of age. Prior to surgery, all animals were group-housed in individual ventilated cages (Optimice, Animal Care Systems, Centennial, CO) on corn-cob bedding (‘Bed-o’-Cobs 1/8 in., The Andersons, Maumee, OH) with a square nestlet and PVC tube provided for enrichment. Three-days prior to surgery, mice were handled daily for approximately 3 minutes and provided with a diet-gel (Dietgel^®^ Recovery, Clear H2O, Westbrook, ME) to allow for acclimatization; continuation of the diet-gel was available for all mice up to 3 days post-operatively. After surgical implantation of a carotid catheter, mice were singly housed to eliminate the risk of catheter port destruction and assessed daily for clinical signs of pain. Mice were allowed free access to rodent chow (LabDiet 5053, St. Louis, MO) and reverse-osmosis-purified water and maintained on a 12:12-h light:dark cycle. Mice were designated as specific pathogen free (SPF) for: mouse rotavirus, Sendai virus, pneumonia virus of mice, mouse hepatitis virus, minute virus of mice, mice parvovirus, Theiler murine encephalomyelitis virus, mouse reovirus type 3, mouse norovirus, lymphocytic choriomeningitis virus, mouse thymic virus, mouse adenovirus types 1 and 2, mouse cytomegalovirus, polyoma virus, K virus, ectromelia virus, Hantavirus, Prospect Hill virus, *Filobacterium rodentium, Encephalitozoon cuniculi*, and *Mycoplasma pulmonis, Helicobacter* spp., *Clostridium piliforme*, and free of any endo- and ectoparasites.

### Experimental design

The experimental workflows of the 2 treatment groups are presented in Figure 1. The SRB patented formulation (SRB-LAB, 1 mg/mL, 5 mL vial) was prepared and acquired from Zoopharm (Windsor, CO). The XRB formulation (Ethiqa XR, 1.3 mg/mL, 3 mL vial) was a proprietary pharmaceutical grade formulation prepared and acquired from Fidelis Pharmaceuticals (North Brunswick, NJ). Beginning 3 days prior to surgery, daily body weights were measured, and mice were clinically evaluated for health status. Mice in each sex grouping were then assigned to either the SRB (n=6 male, n=7 female) or XRB (n=8 male, n=7 female) treatment group. On day 0, surgical catheterization was performed (see next section), and each mouse received either SRB (1 mg/kg, SC, Hamilton syringe, 23g needle) or XRB (3.25 mg/kg, SC, Hamilton syringe, 23g needle) in the right rear flank during the anesthesia induction period. Time at which analgesic was given was recorded and used to determine subsequent blood collection time points. Prior to injection, both SRB and XRB were mixed well by inversion to ensure homogenous suspensions were injected. After surgery, mice were provided with 0.9% sodium chloride (10 mL/kg, SC) in the left rear flank and allowed to recover in a clean cage placed on a heating pad. At 6, 24, 48, and 72-hours after administration of SRB or XRB, arterial blood samples were collected. All mice were humanely euthanized at the 72-hour blood collection timepoint via CO_2_ asphyxiation followed by secondary cervical dislocation.

**Figure 1.**
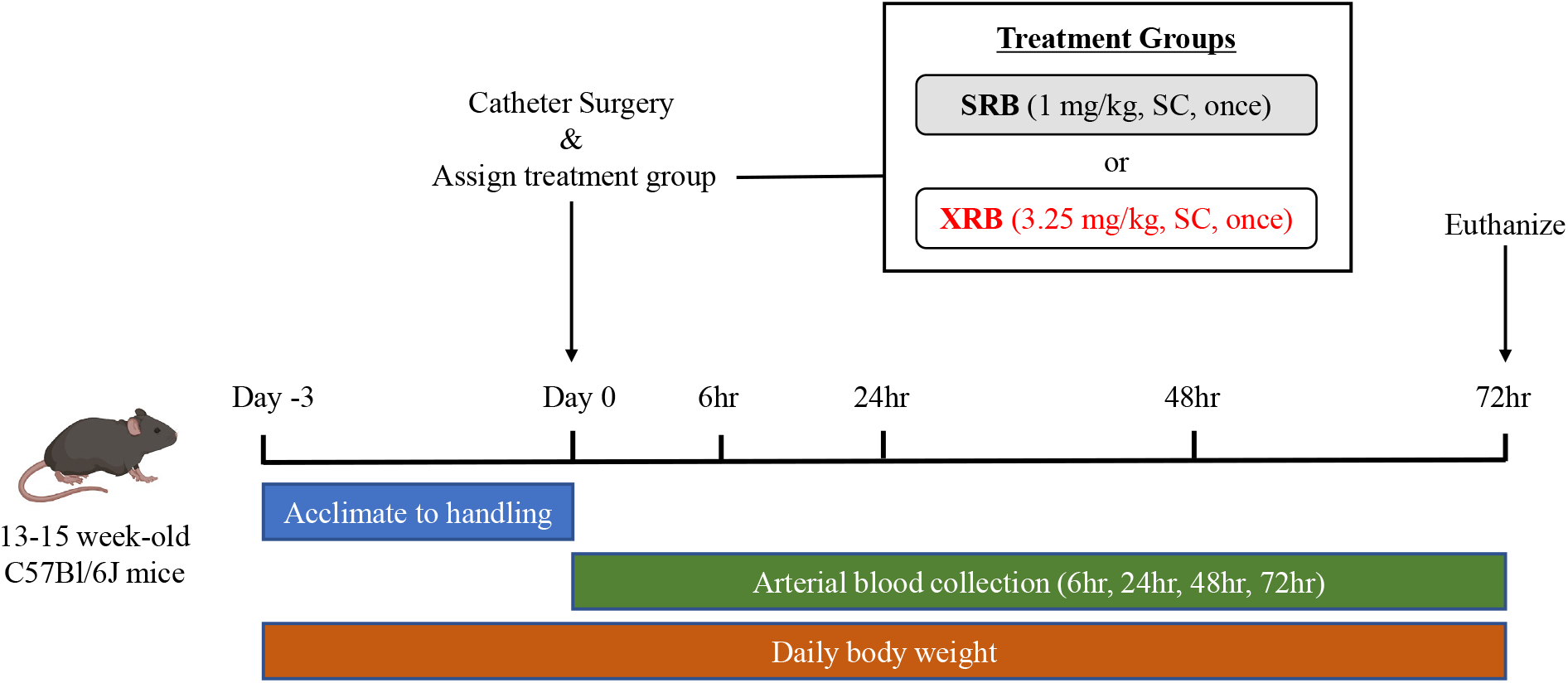
Experimental design. After 3-days of acclimatization, mice were anesthetized, treated with either SRB (1 mg/kg, SC) or XRB (3.25 mg/kg, SC), and surgically implanted with catheters in the left common carotid artery. Arterial blood samples were collected at 4 different time-points (6 hr, 24 hr, 48 hr, and 72 hr) and analyzed for plasma buprenorphine concentrations using LC-MS/MS.

### Surgical procedure

The left carotid artery and right jugular vein were surgically catheterized for blood sampling and infusions, respectively, and the mouse antenna for sampling access^6^ (MASA) was implanted subcutaneously, as previously described in Ayala et al., 2011. Briefly, isoflurane anesthetized mice were aseptically prepared for surgery at the ventral cervical and interscapular regions. A 5-mm longitudinal incision was made just over the left common carotid artery, slightly lateral from the mid-trachea. Using blunt dissection, the common carotid artery was isolated, proximal and distal ligatures were placed, and a silastic-polyethylene −10 catheter^6^ (Fischer Scientific, Waltham, MA), pre-filled with 100-U heparin-saline lock, was secured approximately 10-15 mm into the vessel lumen. The free-end of the catheter was tunneled subcutaneously towards the back of the mouse, exteriorized by a small 5-mm incision in the interscapular region and connected to the MASA device^6^ that sits under the skin. The MASA device was then secured with suture and blood sample ports were easily accessible from the interscapular region of the mouse. The right jugular vein, normally used for infusions, was catheterized, and attached to the MASA in the same fashion as the left carotid artery; however, the infusion side of the MASA was not used in the current study.

### Blood collection

For each blood collection time point, 80 – 100 μL of arterial blood was collected in a blinded fashion from the carotid catheter via the MASA. Care was taken to discard approximately 10 μL of the heparin saline lock flush prior to blood collection, as well as avoidance of introduction of air bubbles into the catheter line. Blood was transferred from a heparin-coated syringe to a 0.5 mL sterile Eppendorf tube (Fisher Scientific, Waltham, MA) on ice and immediately centrifuged at 4 °C and 2500 g for 10 minutes to obtain plasma.^8^ Isolated plasma was transferred to individual sterile glass vials, instantly frozen using liquid nitrogen, and stored at −80 °C until processed for analysis.^13^ After isolation of plasma, saline-washed red blood cells were returned to each mouse via slow infusion into the carotid catheter to avoid anemia. A new 100-U heparin lock was infused into the catheter to maintain patency.

### Pharmacokinetic analysis

Buprenorphine concentrations in mouse plasma were measured using an LC-MS/MS assay established in the City of Hope analytical Pharmacology Core Facility. Following precipitation of plasma proteins with acetonitrile containing 3.5 ng/mL buprenorphine-d4 (Cerilliant, Round Rock, TX, USA) as an internal standard, the sample was vortex mixed for 2 minutes and centrifuged for 10 minutes at 14.8K rpm and 4°C. A 50 μl aliquot of the resulting supernatant was further diluted 1:4 with 40% methanol and 5 μl was injected onto the column. Analyte separation was achieved on a Kinetex 2.6 um C18, 50 x 2.1 mm analytical column (Phenomenex, Torrance, CA USA) using gradient separation. The retention time was 3.6 minutes for buprenorphine and buprenorphine-d4 and the total run time was 7 minutes. Detection was performed using a Xevo TQ-XS Triple Quadrupole Mass Spectrometer (Waters, Milford, MA, USA) with electrospray ionization and operating positive ion mode. The precursor→product ion combinations at m/z 468.38→396.24 for buprenorphine and 472.38→400.24 for buprenorphine-D4 were used in multiple reaction monitoring (MRM) mode, and MassLynx version 4.2 software was used to acquire and analyze data. The lower limit of detection for buprenorphine concentrations was 0.3 ng/mL.^22^

PK parameters were calculated using PK Solver 2.0 with a noncompartmental analysis (NCA) linear up-log down method.^34^ Parameters included in the analysis were half-life, time to maximum concentrations, peak concentrations, area under the concentration-time curve, and clearance (t _1/2_, T _max_, C _max_, AUC _0-last_, CL). For statistical analysis, a two-way ANOVA followed by Bonferroni multiple comparisons test was used to compare differences in buprenorphine concentrations over the 6, 24, 48, and 72-hour sample period. All statistical analyses were complete using GraphPad Prism version 9.2.0 for Windows (San Diego, CA).

## Results

### Overall animal health

During the acclimation period, all animals were handled for approximately 3 minutes daily and had no significant changes in body weight during the preoperative period. Both SRB- and XRB-treated mice had similar declines in body weight after surgical implantation of the catheters. As expected, body weights declined from 1-3 days and 2-3 days post-operatively in male and female mice, respectively (Figure 2A). However, no significant difference in body weight changes were observed between SRB- or XRB-treated mice (Figure 2B). Compared to pre-operative measures, body weights declined by 11.7 ± 1.6% and 12.3 ± 0.7% in males and 7.6 ± 2.2% and 8.1 ± 1.1% (mean ± SEM) in females treated with SRB and XRB, respectively (Figure 2B). For both treatment groups, there were no adverse reactions at the subcutaneous injection site or signs of infection at the catheter implantation site.

**Figure 2.**
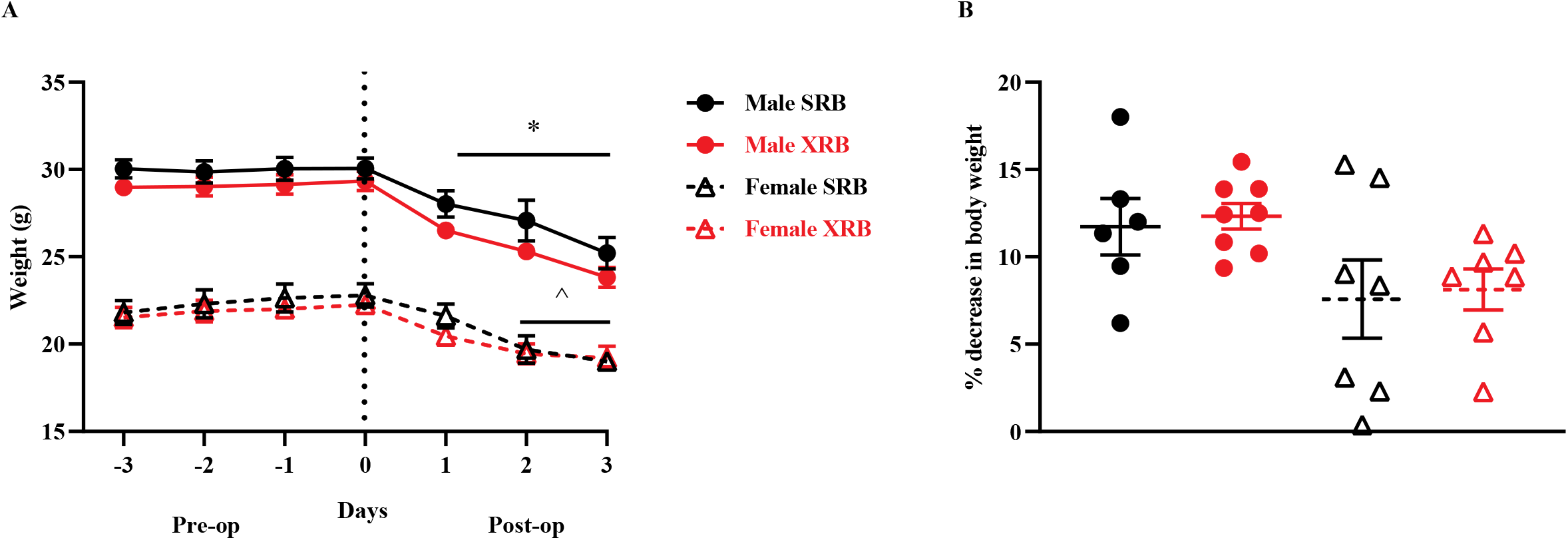
As expected, all mice lost body weight post-operatively, with similar declines in body weight between XRB- and SRB-treated animals. Body weights of XRB- and SRB-treated mice declined significantly 1-3 days (*) and 2-3 days (^) post-operatively in males and females, respectively, but not between XRB- or SRB-treatment groups (A). Percent decreases in body weight were similar between sexes treated with SRB or XRB (B). Data are presented as mean ± SEM, *p-value < 0.05*. Dashed line in (A) indicates day of surgery.

### Pharmacokinetics

Whole blood withdrawn from carotid arterial catheters was assessed for plasma buprenorphine concentrations at 6, 24, 48, and 72-hours after administration of SRB or XRB (Figure 3). At 6, 24, 48, and 72 hours, respective circulating concentrations of buprenorphine (ng/mL) were (mean ± SEM) 3.76 ± 0.55, 2.45 ± 0.36, 1.35 ± 0.17, and 1.02 ± 0.21 for SRB, and 13.52 ± 1.89, 7.37 ± 1.20, 4.40 ± 0.69, and 3.23 ± 1.0 for XRB. XRB circulating buprenorphine concentrations were significantly greater than SRB concentrations at 6, 24, and 48-hours (p < 0.05), but not at 72-hours after administration. In surgically catheterized mice, circulating buprenorphine concentrations remained above the therapeutic threshold of 1 ng/mL^19,22^ (dashed line, Figure 3) for 48 – 72 hours after administration of SRB and XRB. According to the NCA, the slope of elimination for plasma buprenorphine concentration over time shows an exponential decline for SRB (R^2^ = 0.98) and XRB (R^2^ = 0.96) based on the following equations:

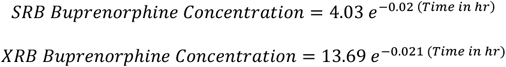

**Figure 3.**
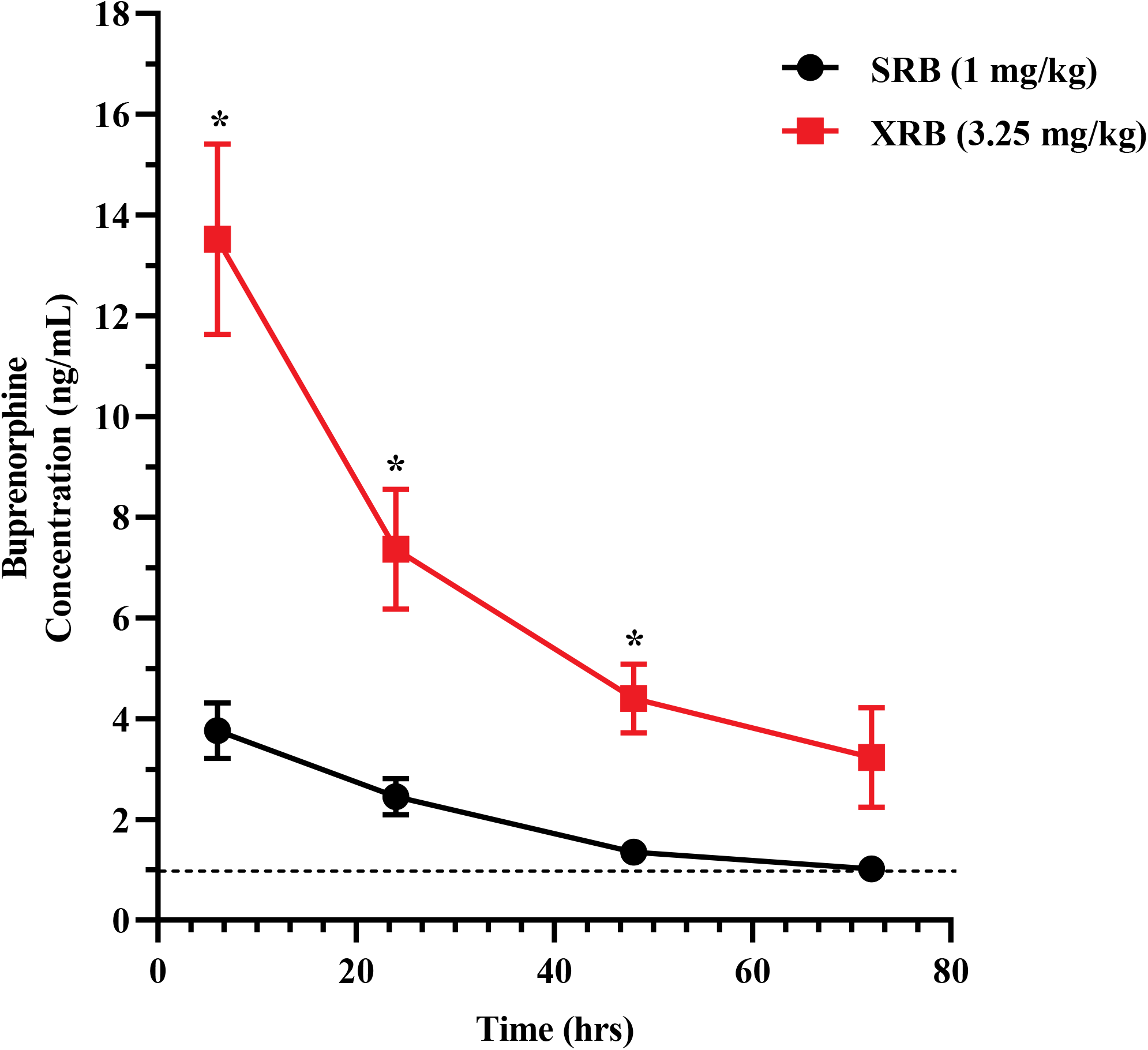
Mice treated with SRB (1 mg/kg, SC, black line) or XRB (3.25 mg/kg, SC, red line) had circulating buprenorphine levels that were above the therapeutic effective threshold of 1 ng/mL^19,33^ (dashed line) up to 72 hours post-operatively. Data are presented as mean ± SEM, * = *p-value < 0.05*.

Pharmacokinetic parameters for SRB- and XRB-treated mice are described in Table 1. Half-life (T_1/2_) and clearance (Cl) were similar between mice treated with SRB or XRB. Although both SRB and XRB had maximum plasma buprenorphine concentrations 6-hours after administration, maximum buprenorphine concentrations (C_max_, ng/mL) were three-to-four times greater in XRB- than SRB-treated mice. Additionally, area under the curve (AUC_0-last_), a measure typically associated with systemic exposure of a drug, was greater in XRB- than SRB-treated animals.

**Table 1.** Pharmacokinetic Parameters of SRB and XRB were calculated from a non-compartmental pharmacokinetic analysis^34^ (NCA) of averaged buprenorphine concentrations at each time point (6 hr, 24 hr, 48 hr, and 72 hr). t _1/2_ = half-life, T _max_ = time to maximum concentrations, C _max_ = peak concentration, AUC _0-last_ = area under the concentration-time curve, and CL = clearance.

| Parameter | Unit | SRB | XRB |
| --- | --- | --- | --- |
| $t_{1/2}$ | h | 37.82 | 40.32 |
| $T_{max}$ | h | 6 | 6 |
| $C_{max}$ | ng/mL | 3.76 | 13.52 |
| $AUC_{0-last}$ | (ng/mL) × h | 138.98 | 451.90 |
| Clearance | μL / h / kg | 5.14 | 5.08 |

## Discussion

In this study, we surgically implanted indwelling carotid arterial and jugular vein catheters and assessed the pharmacokinetic response of subcutaneously administered SRB and XRB. This study design enabled comparison of plasma buprenorphine concentrations up to 72-hours after administration of SRB or XRB. We identified that both SRB and XRB had similar half-lives and maintained circulating buprenorphine concentrations above the therapeutic level of 1.0 ng/mL for 72-hours after administration. Furthermore, we determined that the maximum concentration and AUC of plasma buprenorphine was three-to-four times greater in XRB than SRB-treated mice, likely due to the approximate three-fold increase in the recommended dose of XRB (3.25 mg/kg) compared to SRB (1 mg/kg). These results support our hypothesis that post-operatively, mice treated with either SRB or XRB would have circulating buprenorphine concentrations exceeding the therapeutic threshold for up to 72-hours.

The goal of this study was to directly compare the pharmacokinetic parameters of the newly available XRB with the commonly used SRB. In humans, buprenorphine concentrations of at least 1.0 ng/mL provided pain relief in clinical reports^10,31^; thus, buprenorphine concentrations of at least 1.0 ng/mL in rodents have been accepted as a therapeutic threshold providing adequate pain relief.^19,22^ As previously reported, we confirmed that SRB-treated mice had circulating buprenorphine concentrations above 1.0 ng/mL for 48-72-hours after administration; although, the 72-hour buprenorphine concentrations teetered along the borderline threshold level.^21^ XRB-treated mice maintained buprenorphine concentrations above 1.0 ng/mL for up to 72-hours. (Figure 3). These results establish that XRB is comparable to SRB in providing effective therapeutic buprenorphine concentrations in mice for the alleviation of post-surgical pain.

Alleviation of post-surgical pain in mice is commonly accomplished using different formulations of long-lasting buprenorphine; however, higher doses of these formulations have in some cases resulted in respiratory depression, inflammatory changes, subcutaneous hemorrhage, and necrosis at the site of injection.^9,20^ In rats, a subcutaneous lipid suspension of buprenorphine at a dose of 6.25 mg/kg resulted in inflammatory changes and hemorrhage at or near the injection site.^20^ One study in mice treated with SRB at a dose of 1 mg/kg revealed scabby lesions with local erythema at the injection site.^9^ The authors attributed this to the seeping out of SRB onto the skin. A recent study of PK and histopathological lesions in rats administered XRB revealed mild-to-moderate granulomatous changes at injection sites at both low- and high-dosages (0.65 mg/kg and 1.30 mg/kg, respectively).^25^ In the current study, although histopathological lesions at injection site were not assessed, we did not observe any clinical or gross adverse reaction of SRB or XRB at the injection site. Furthermore, we used a Hamilton syringe, injected slowly, and pinched the skin after administration to ensure the drugs would not seep out. We recommend that others use this delivery method with both SRB and XRB to avoid any inadvertent skin reactions.

According to the *NIH-ARAC Guidelines for Blood Collection in Mice and Rats*, the amount of blood that can be safely withdrawn from a single mouse is approximately 10% of the total blood volume (72 ml/kg, mean) every 2 to 4 weeks, 7.5% every 7 days, and 1% every 24 hours^3^. These guidelines are important to prevent anemia, dehydration, and associated pain and distress. However, PK studies often require larger amounts of blood, depending on the compound in question, with 6-12 terminal blood samples per compound per time point necessitating higher animal usage and increased variability between samples^24^. In the current study, the indwelling carotid catheter enabled us to collect multiple blood samples over a 72-hour period, return saline-washed red blood cells to each mouse, and assess buprenorphine concentrations from the same mouse at different time points. This method is an important refinement to the commonly used PK mouse design paradigms, where larger volumes of blood are collected from only one mouse per time point.^14,29^ Studies have shown varied responses of injectable buprenorphine based on mouse strain and individual animals; thus, the use of an indwelling catheter minimizes intra-animal variation and potentially increases translatability.^6,7,29^ However, for PK studies requiring larger volumes of blood (i.e. greater than 10% of the total blood volume) and where returning washed red blood cells is not optimal, indwelling catheterization of mice will likely be inadequate and larger animal models, such as rats, should be used.^14^ Importantly, assessment of PK parameters acquired from the same animal over time more closely mimics preclinical experimental drug trials that are performed in humans and larger animal species.^14,29^ Therefore, the use of an indwelling carotid catheter for pharmacokinetic blood collection in mice may be superior for studies requiring multiple blood collections in volumes up to 100 μl per mouse.

There are several differences between SRB and XRB that may influence when to choose one over the other. SRB is a USP grade patented formulation containing a solvent with N-methyl-2-pyrrolidone. Administration of SRB in mice is considered extra-label use; however, at adequate dosages many studies have shown its safety and efficacy in multiple strains of mice.^4,9,12,17,21,22^ XRB, on the other hand, is an FDA-indexed cGMP formulation that uses a lipid-based delivery vehicle.^19,20,30^ FDA-indexed drugs go through an alternative FDA approval process for species that are “too rare or varied” to undergo a full approval process. FDA-indexing of drugs takes less time than the standard FDA approval process and decreases expenses without compromising the efficacy of the drug or the safety of the animals; thus, XRB has a level of FDA-certification that is currently lacking in SRB.^5^ Yet, SRB has many third party studies ensuring both its safety efficacy in the rodent species.^9,11,12,17,21,22^ Nonetheless, both SRB and XRB meet standards set by *The Guide* for ensuring “toxic or unwanted side effects” are not associated with drug administration.^2^ Our results support the use of either SRB or XRB for alleviation of post-surgical pain in mice and the new availability of XRB increases the options for safe and effective analgesia in rodent models.

## Abbreviations and Acronyms

SRB: sustained-release buprenorphine
XRB: extended-release buprenorphine
MASA: mouse antenna for sampling access
NCA: non-compartmental analysis

## Acknowledgements

The authors thank the City of Hope Analytical Pharmacology and Animal Resources Cores supported by the National Cancer Institute of the NIH (P30CA033572). We thank the staff of the Comprehensive Metabolic Phenotype Core & Center for Comparative Medicine. Fidelis Pharmaceuticals provided the XRB used in this study. M.S. was supported by a National Cancer Institute Cancer Metabolism Training Program Postdoctoral Fellowship (T32CA221709). This work was supported, in part, by intramural funding from the Beckman Research Institute of City of Hope.

## References

1. [Internet]. AALAS position statement on Alleviating pain and distress in laboratory animals. Available at: https://www.aalas.org/about-aalas/position-papers/alleviating-pain-and-distress.

2. 2011. Guide for the Care and Use of Laboratory Animals, eigth edition 8th ed. Washington (DC): The National Academic Press.

3. [online] N-AG. Guidelines for Blood Collection in Mice and Rats. NIH.

4. Adamson TW, Kendall LV, Goss S, Grayson K, Touma C, Palme R, Chen JQ, Borowsky AD. 2010. Assessment of carprofen and buprenorphine on recovery of mice after surgical removal of the mammary fat pad. J Am Assoc Lab Anim Sci 49:610–616.

5. Adminsitration USFaD. [Internet]. 2021. Drug Indexing Available at: https://www.fda.gov/animal-veterinary/minor-useminor-species/drug-indexing.

6. Ayala JE, Bracy DP, Malabanan C, James FD, Ansari T, Fueger PT, McGuinness OP, Wasserman DH. 2011. Hyperinsulinemic-euglycemic Clamps in Conscious, Unrestrained Mice. JoVE:e3188.

7. Bardelmeijer HA, Buckle T, Ouwehand M, Beijnen JH, Schellens JH, van Tellingen O. 2003. Cannulation of the jugular vein in mice: a method for serial withdrawal of blood samples. Lab Anim 37:181–187.

8. Blankenship-Paris TL, Dutton JW, Goulding DR, McGee CA, Kissling GE, Myers PH. 2016. Evaluation of buprenorphine hydrochloride Pluronic® gel formulation in male C57BL/6NCrl mice. Lab Animal 45:370–379.

9. Carbone ET, Lindstrom KE, Diep S, Carbone L. 2012. Duration of action of sustained-release buprenorphine in 2 strains of mice. J Am Assoc Lab Anim Sci 51:815–819.

10. Chawarski MC, Schottenfeld RS, O’Connor PG, Pakes J. 1999. Plasma concentrations of buprenorphine 24 to 72 hours after dosing. Drug Alcohol Depend 55:157–163.

11. Chum HH, Jampachairsri K, McKeon GP, Yeomans DC, Pacharinsak C, Felt SA. 2014. Antinociceptive effects of sustained-release buprenorphine in a model of incisional pain in rats (Rattus norvegicus). J Am Assoc Lab Anim Sci 53:193–197.

12. Clark TS, Clark DD, Hoyt RF, Jr. 2014. Pharmacokinetic comparison of sustained-release and standard buprenorphine in mice. J Am Assoc Lab Anim Sci 53:387–391.

13. DenHerder JM, Reed RL, Sargent JL, Bobe G, Stevens JF, Diggs HE. 2017. Effects of Time and Storage Conditions on the Chemical and Microbiologic Stability of Diluted Buprenorphine for Injection. J Am Assoc Lab Anim Sci 56:457–461.

14. Diehl KH, Hull R, Morton D, Pfister R, Rabemampianina Y, Smith D, Vidal JM, van de Vorstenbosch C. 2001. A good practice guide to the administration of substances and removal of blood, including routes and volumes. J Appl Toxicol 21:15–23.

15. Falkenberg MK, Teilmann AC, Henriksen T, Hau J, Poulsen HE, Abelson KS. 2019. Clinical, Physiologic, and Behavioral Evaluation of Permanently Catheterized NMRI Mice. J Am Assoc Lab Anim Sci 58:380–389.

16. Foley PL, Kendall LV, Turner PV. 2019. Clinical Management of Pain in Rodents. Comp Med 69:468–489.

17. Foley PL, Liang H, Crichlow AR. 2011. Evaluation of a sustained-release formulation of buprenorphine for analgesia in rats. J Am Assoc Lab Anim Sci 50:198–204.

18. Gades NM, Danneman PJ, Wixson SK, Tolley EA. 2000. The magnitude and duration of the analgesic effect of morphine, butorphanol, and buprenorphine in rats and mice. Contemporary topics in laboratory animal science 39:8–13.

19. Guarnieri M, Brayton C, DeTolla L, Forbes-McBean N, Sarabia-Estrada R, Zadnik P. 2012. Safety and efficacy of buprenorphine for analgesia in laboratory mice and rats. Lab Anim (NY) 41:337–343.

20. Guarnieri M, Brayton C, Sarabia-Estrada R, Tyler B, McKnight P, DeTolla L. 2017. Subcutaneous Implants of a Cholesterol-Triglyceride-Buprenorphine Suspension in Rats. Journal of veterinary medicine 2017:3102567.

21. Healy JR, Tonkin JL, Kamarec SR, Saludes MA, Ibrahim SY, Matsumoto RR, Wimsatt JH. 2014. Evaluation of an improved sustained-release buprenorphine formulation for use in mice. American journal of veterinary research 75:619–625.

22. Kendall LV, Hansen RJ, Dorsey K, Kang S, Lunghofer PJ, Gustafson DL. 2014. Pharmacokinetics of sustained-release analgesics in mice. J Am Assoc Lab Anim Sci 53:478–484.

23. Lascelles BDX, Waterman AE, Cripps PJ, Livingston A, Henderson G. 1995. Central sensitization as a result of surgical pain: investigation of the pre-emptive value of pethidine for ovariohysterectomy in the rat. PAIN 62.

24. Leblanc AF, Huang KM, Uddin ME, Anderson JT, Chen M, Hu S. 2018. Murine Pharmacokinetic Studies. Bio Protoc 8:e3056.

25. Levinson BL, Leary SL, Bassett BJ, Cook CJ, Gorman GS, Coward LU. 2021. Pharmacokinetic and Histopathologic Study of an Extended-Release, Injectable Formulation of Buprenorphine in Sprague-Dawley Rats. J Am Assoc Lab Anim Sci 60:462–469.

26. Medina LV, Hrapkiewicz K, Tear M, Anderson LC. 2007. Fundamental training for individuals involved in the care and use of laboratory animals: a review and update of the 1991 NRC Core Training Module. Ilar j 48:96–108.

27. Pacharinsak C, Smith JC. 2017. Handbook of laboratory animal anesthesia and pain management: rodents. Boca Raton: CRC Press, Taylor & Francis Group.

28. Penderis J, Franklin RJM. 2005. Effects of pre-versus post-anaesthetic buprenorphine on propofol-anaesthetized rats. Veterinary Anaesthesia and Analgesia 32:256–260.

29. Peng SX, Rockafellow BA, Skedzielewski TM, Huebert ND, Hageman W. 2009. Improved pharmacokinetic and bioavailability support of drug discovery using serial blood sampling in mice. J Pharm Sci 98:1877–1884.

30. Traul KA, Romero JB, Brayton C, DeTolla L, Forbes-McBean N, Halquist MS, Karnes HT, Sarabia-Estrada R, Tomlinson MJ, Tyler BM, Ye X, Zadnik P, Guarnieri M. 2015. Safety studies of post-surgical buprenorphine therapy for mice. Lab Anim 49:100–110.

31. Watson PJ, McQuay HJ, Bullingham RE, Allen MC, Moore RA. 1982. Single-dose comparison of buprenorphine 0.3 and 0.6 mg i.v. given after operation: clinical effects and plasma concentration. Br J Anaesth 54:37–43.

32. Woolf CJ, Wall PD. 1986. Morphine-sensitive and morphine-insensitive actions of C-fibre input on the rat spinal cord. Neurosci Lett 64:221–225.

33. Yassen A, Olofsen E, Dahan A, Danhof M. 2005. Pharmacokinetic-pharmacodynamic modeling of the antinociceptive effect of buprenorphine and fentanyl in rats: role of receptor equilibration kinetics. The Journal of pharmacology and experimental therapeutics 313:1136–1149.

34. Zhang S, Baker J, Pulay P. 2010. A reliable and efficient first principles-based method for predicting pK(a) values. 2. Organic acids. J Phys Chem A 114:432–442.

35. ZooPharm. Windsor, CO: ZooPharm.

